# Scalable summary statistics-based heritability estimation method with individual genotype level accuracy

**DOI:** 10.1101/2024.03.09.584258

**Authors:** Moonseong Jeong, Ali Pazokitoroudi, Zhengtong Liu, Sriram Sankararaman

## Abstract

SNP heritability, the proportion of phenotypic variation explained by genotyped SNPs, is an important parameter in understanding the genetic architecture underlying various diseases and traits. Methods that aim to estimate SNP heritability from individual genotype and phenotype data are limited by their ability to scale to Biobank-scale datasets and by the restrictions in access to individual-level data. These limitations have motivated the development of methods that only require summary statistics. While the availability of publicly accessible summary statistics makes them widely applicable, these methods lack the accuracy of methods that utilize individual genotypes.

Here we present a SUMmary statistics-based Randomized Haseman-Elston regression (SUM-RHE), a method that can estimate the SNP heritability of complex phenotypes with accuracies comparable to approaches that require individual genotypes, while exclusively relying on summary statistics. SUM-RHE employs Genome-Wide Association Study (GWAS) summary statistics and statistics obtained on a reference population, which can be efficiently estimated and readily shared for public use. Our results demonstrate that SUM-RHE obtains estimates of SNP heritability that are substantially more accurate compared to other summary statistic methods and on par with methods that rely on individual-level data.

## 1 Introduction

The exponentially decreasing cost of genotyping and sequencing technologies has led to an increase in the number and size of biobanks ^1,2,3^, covering a wide range of populations. With large samples of phenotype and genotype data now available in these biobanks, one of the major analyses often performed is estimating heritability, defined as the phenotypic variance explained by the variance in the genotype ^4^. Heritability estimates in these large data sets have provided new insights into the underlying genetic architecture of complex traits ranging from schizophrenia ^5^ to height^6^. Most methods fit linear mixed models (LMMs) ^7,8,9^ to map the variation in genotypes measured at single nucleotide polymorphisms (SNPs) to the variation in phenotypes and thereby estimate the SNP heritability, *i*.*e*., the proportion of phenotypic variance explained by genotyped SNPs. Given the high dimensionality of the genotypes and the large sample sizes of biobanks, fitting or parameter estimation in LMMs is computationally prohibitive. Many methods have been proposed to reduce computational complexity while retaining statistical accuracy ^7,8,9,10,11,12,13^. These methods, while highly accurate, generally take hours or days to run and require access to individual genotypes and phenotypes.

The rise of large-scale biobanks has also brought increased attention to the issue of genomic privacy due to a surge in security breaches. Consequently, additional measures have been implemented to safeguard individual information throughout its processing, storage, and sharing ^14^. Gaining access to raw individual-level data is now more challenging, exemplified by the UK Biobank’s decision to restrict access to its WGS data to its cloud server, resulting in added server costs for analysis. Given this development, there is a growing preference for summary statistics-based methods due to their portability and speed, even though they may sacrifice some statistical power compared to methods that use individual-level data ^10,15,16,17,18^. Such a loss in statistical power is particularly pronounced in smaller sample sizes, and may result in inflated estimates of heritability due to underestimation of linkage disequilibrium (LD) ^10^, even if correct reference summary statistics were used.

To address these challenges of heritability estimation in large biobanks, we propose SUM-RHE (SUMmary-statistics Randomized Haseman-Elston regression), by extending our previous work, Randomized Haseman-Elston regression (RHE) ^11,12^, to work exclusively on summary statistics. This adaptation leverages the observation that the trace estimates of the squared genetic relatedness matrix (GRM), that are needed to compute the method-of-moments (MoM) estimator underlying RHE, can be related to population-level parameters. By combining these trace estimates from a reference sample with GWAS summary statistics from a target sample (consisting of individuals sampled from the same population as the reference sample), we can reconstruct the MoM estimates for the target sample without access to the individual data. In comprehensive simulations across various genetic architectures and scenarios, we show that SUM-RHE estimates are on par with methods that rely on individual-level data and substantially more accurate than summary statistic-based methods, all while exclusively utilizing summary statistics.

## 2 Method

### 2.1 Background on heritability estimation and linear mixed models

Early attempts to calculate SNP heritability of complex traits by aggregating SNPs identified as GWAS significant have revealed the issue of missing heritability, as this estimate of heritability was significantly lower than the narrow-sense heritability estimated in other studies (e.g., twin studies ^19^). The seminal work by Yang *et al*. ^20^ reduced this discrepancy by jointly modeling all the SNPs, such that their effect sizes come from a distribution of some fixed variance that quantifies the genetic variation. In this LMM framework, the standardized phenotype vector ***y*** is modeled as a linear combination of SNP effect sizes ***β*** multiplied by the standardized genotype matrix ***X*** of *M* SNPs and *N* individuals with uniform noise ***ϵ***:

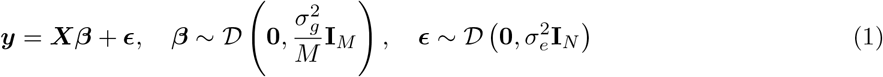

Where the additive effect sizes ***β*** are drawn from an arbitrary distribution *𝒟* with mean zero and variance of 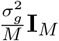, and the environmental/noise effects ***ϵ*** drawn from a distribution with variance 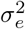. In the original work by Yang *et al*. ^20^ and GCTA^7^, the distribution *𝒟* was chosen as a Normal distribution. The SNP-heritability is then defined as the proportion of genetic variance over total phenotypic variance, 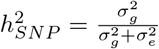.

One approach to estimating the variance components 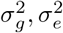 is to find the maximum likelihood estimator (MLE) and variants such as restricted maximum likelihood (REML) estimators ^7,8^. These methods often rely on iterative optimizations, which tend to be inefficient, and could lead to biased estimates due to the normality assumption ^10,12^. On the other hand, MoM approaches such as the Haseman-Elston regression (HE) ^21,22^, randomized Haseman-Elston regression (RHE) ^12,11^, MQS ^10^, or LDSC^15^, only require solving the normal equations and do not make any assumptions on the underlying distribution *𝒟*. Here, we briefly discuss the method-of-moments estimator (HE regression), which sets the foundation for our work.

### 2.2 Heritability estimation from individual genotype data using method of moments and randomized method of moments

The HE MoM estimator of the parameters 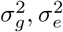, can be obtained by minimizing the discrepancy between the population covariance and the sample covariance matrices ***yy***^T^. The population covariance is given as:

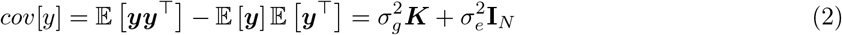

Where 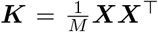 is defined as the genetic relatedness matrix (GRM). We want to find the estimates of the parameters 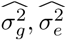 that minimize the Frobenius norm (the measure of discrepancy) between the two covariance matrices. This is equivalent to solving the normal equations:

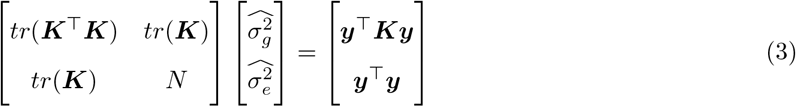

Where *tr*(***K***) = *N* and ***y***^T^***y*** = *N*, given both ***X*** and ***y*** are standardized. Equation (3) has the analytical solution for the variance components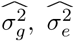 :

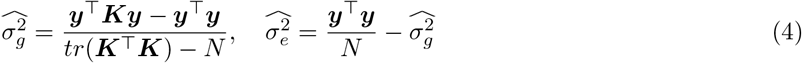

Giving the MoM estimate for heritability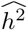 :

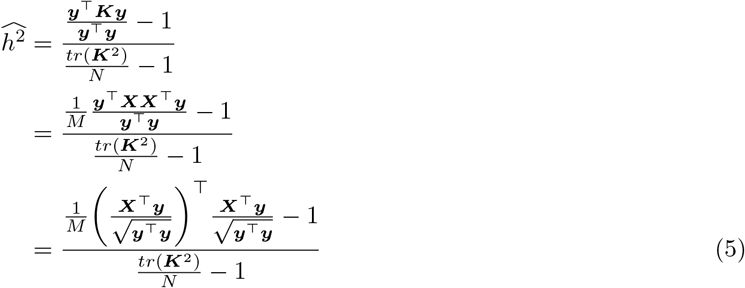

The biggest bottleneck in the equation is calculating *tr* (***K***^T^***K*)**. An exact calculation of the trace involves forming the matrix ***K***^T^***K*** which has a computational complexity of *𝒪*(*MN*^2^). Given *M ≈* 1, 000, 000 and *N ≈* 1, 000, 000, this is not tractable in modern biobanks. One of the main contributions of RHE-mc ^11^ and RHE-reg ^12^ is the efficient estimation of *tr (****K***^T^***K)*** by leveraging the fact that the trace of ***K***^T^***K*** can be approximated by a stochastic trace estimator ^23,24^:

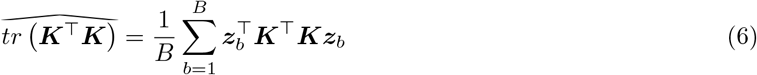

Where ***z***_*b*_ are i.i.d random vectors such that 𝔼 [***z***_*b*_] = **0** and 𝔼 ***z***_*b*_***z***_*b*_T = **I**_*N*_. In both RHE-mc and RHE-reg, random vectors sampled from the standard normal distribution are used. Numerically, it was found that using *B ≈* 100 can estimate the trace of the squared GRM matrix ***K***^T^***K*** with high accuracy for moderate sample sizes of *N ≈* 5000 and *B ≈* 10 for large samples sizes ^11,12^. This stochastic trace estimator reduces the computational complexity to *𝒪* (*MNB*). Additional optimizations, such as the mailman algorithm ^25^ and implementation of a streaming version of the algorithm, reduce the computational complexity to 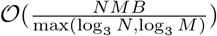 and the memory complexity to *𝒪* (*NB*), thus allowing estimation of heritability across millions of SNPs and individuals.

Extensive benchmarking of the RHE methods has shown that their performance is on par with other methods that require individual-level data, such as GCTA or BOLT-REML^11,12^. RHE offers a distinct advantage over these likelihood-based methods, which tend to scale poorly, as well as other MoM approaches in terms of computational and memory efficiency (e.g., HE) or statistical efficiency (LDSC) ^11^. However, RHE is still limited in that it requires access to individual-level genotype and phenotype data, which restricts its applicability in cases where such data is unavailable or only GWAS summary statistics are available.

### 2.3 Heritability estimation from summary statistics

In this work, we further extend RHE to work exclusively with GWAS summary statistics. Our key observation is the fact that the left-hand side (LHS) of the normal equations (Equation 3) is related to the LD in the population ^10,26^ and not on the phenotype. Thus, if we can summarize the trace estimate for a reference sample drawn from a population, we can use these trace estimates to recreate the corresponding trace estimates in a target sample drawn from the same population. Indeed, we find that the expected value of the trace of ***K***^T^***K*** can be related to the LD scores of the SNPs. Further, the RHS of the normal equations can be computed from GWAS summary statistics obtained on the target population.

The LD score of a variant *j* is defined as the sum of squared correlation with all the variants:

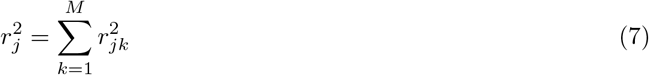

Then *tr*(***K***^T^***K***) is:

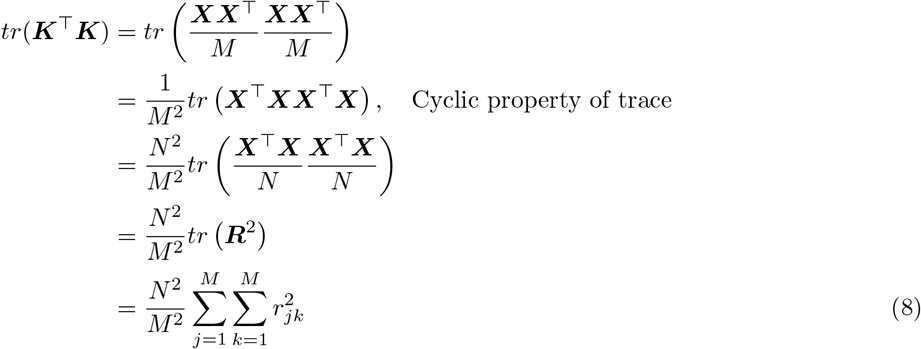

Where ***R*** is the *M × M* correlation matrix (or the LD matrix) and 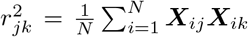 is the sample correlation between variants *j, k*:

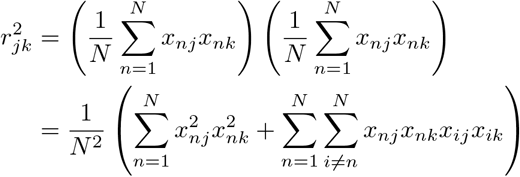

For large sample sizes (*N*), we have

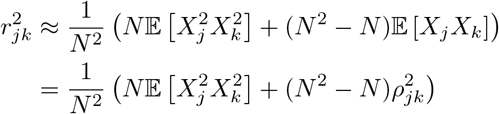

Here *ρ*_*jk*_ is the expected correlation or population LD between SNPs *j* and *k*. Assuming (*X*_*j*_, *X*_*k*_) are normally distributed with mean zero and covariance 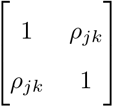, we can use Isserlis’ theorem to compute 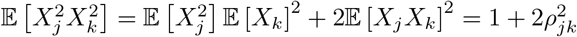. We then have:

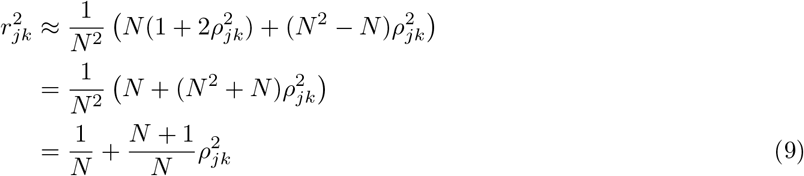

Substituting Equation 9 into Equation 8,

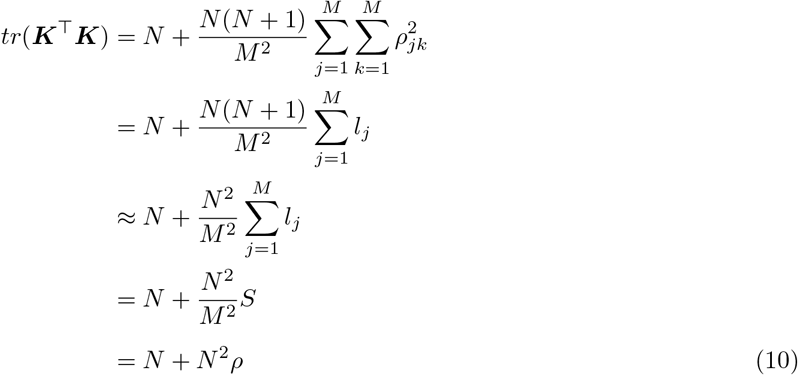

Here 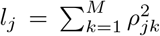 is the expected or population-level LD score associated with SNP *j* while *ρ* can be interpreted as the average (expected) LD across all SNPs in genotype ***X***.

Let 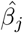 denote the GWAS effect size estimates for SNP *j* obtained by linear regression. Given the observed count *N*_*j*_ of the SNP, we have 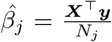 since the genotypes are standardized. The standard error of the GWAS estimate,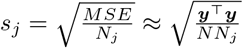. Thus, if we define the vector of adjusted z-scores:

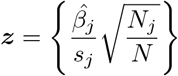

We have that

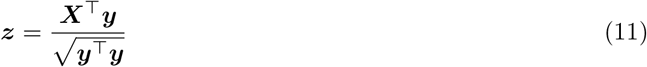

Substituting Equations 10 and 11 into Equation 5 gives us:

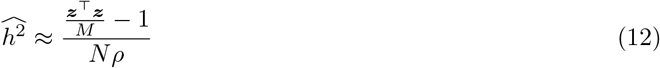

Here.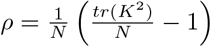 Given 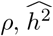 can be computed using summary statisics using Equation 12. However, computing *ρ* requires the exact *κ* = *tr*(*K*^2^), which is computationally intractable. Instead we approximate *ρ* by using the stochastic trace estimates 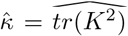 described in Equation 6, such that 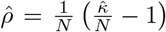 This gives us the randomized Method of Moments estimator of *h*^2^ that can be calculated from summary statistics:

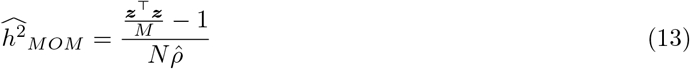

We propose releasing 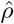 as “trace summaries”, which can then be combined with phenotype-specific GWAS summary statistics ***z*** to estimate heritability. The estimator in Equation 14 assumes that the 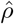 was computed on the same genotypes used to generate the GWAS summary statistics. In settings where 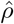 cannot be computed on the same genotypes, we can use 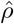 computed on a reference genotype dataset drawn from a population that is similar to the population that was used to generate GWAS summary statistics (such as the 1000 Genomes project). This is under the assumption that the LD structures of similar populations will also be related.

#### 2.3.1 Estimating standard errors

To calculate the standard error of our estimator, we perform SNP-level block jackknife resampling, as done in RHE-mc ^11^. When generating the trace summaries with individual genotypes, we report the 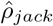 values estimated from jackknife subsamples. Excluding the same SNPs from the PLINK GWAS summary statistics, we can compute the denominator in Equation 14, 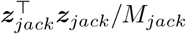, to get the jackknife replicate of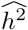 :

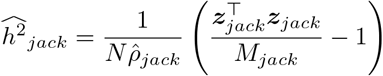

Following the execution on all SNP blocks, we employ jackknife resampling to obtain SE estimates:

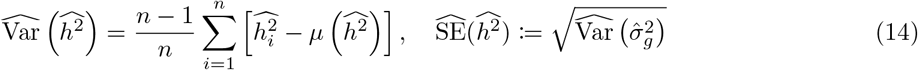

## 3 Results

### 3.1 Simulations under varied genetic architectures

We assessed the performance of SUM-RHE against methods that require individual genotypes: RHE (RHE-mc run with a single component), GCTA-GREML, and BOLT-REML, and methods that can work with summary statistics: LDSC and SumHer ^16^, on the task of estimating genome-wide heritability. We applied all methods to unrelated white British individuals genotyped on *M* = 454, 207 common SNPs (MAF T 0.01 excluding SNPs in the MHC region) typed on the UKBiobank Axiom array. Due to the computational scalability of GCTA-GREML and BOLT-REML, we tested these and other methods in a small-scale setting where the number of individuals in the target dataset was set to *N*_*target*_ = 10, 060 (we term this the 10*k* sample). In addition, we compared all the remaining methods in a large-scale setting where the number of individuals in the target dataset was set to *N*_*target*_ = 50, 112 (termed the 50*k* sample). For the summary statistic methods, the GWAS summary statistics were computed on the target datasets. These methods also require population statistics, calculated in the remainder of the *N* = 291, 273 unrelated white British individuals as the reference dataset. For the small-scale simulation, the reference set has *N*_*ref*_ = 281, 213, and for the large-scale simulation, we set *N*_*ref*_ = 241, 161. On each of the reference sets, we generated SUM-RHE trace summary statistics, LDSC reference LD scores, and LDAK SNP taggings.

We then simulated phenotypes corresponding to 9 different genetic architectures: 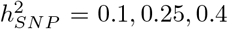 and causal ratio *p* = 1.0, 0.1, 0.01 (where the causal ratio represents the proportion of variants with non-zero effects), each with 100 replicates. Table 1 below summarizes the inputs for each method: For the calculation of LDSC LD scores, we used the entire *N*_*ref*_ reference sample with a window size of 2Mb. SUM-RHE trace summaries were calculated by aggregating the trace estimates of 25 runs on the reference set with *B* = 100 (equivalent to stochastic trace estimation with *B*^*′*^ = 2500 random vectors; see Figure S2 for more information) and 1000 jackknife blocks, yielding a single trace summary statistic with 1000 jackknife estimates of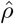. SumHer was run assuming the GCTA model to calculate the SNP taggings (consistent with the genetic architecture assumed in our simulations). RHE was run with *B* = 100 and 1000 jackknife blocks as well. BOLT-REML/GCTA-GREML were run with default parameter settings. GWAS summary statistics for each simulated phenotype were generated using PLINK 2.0. Figure 1 below summarizes the heritability estimates on the target data. Across the 18 different settings (genetic architectures and sample sizes) we tested, we found that the accuracy of SUM-RHE was comparable to RHE (Figure 1) with the mean-squared error of SUM-RHE close to 1 relative to RHE, despite relying only on the summary statistics (Figure 2; see Figure S1 for the MSE of each method).

**Table 1:** Inputs for the different methods evaluated. LDSC and SUM-RHE rely only on the summary statistics, while GCTA-GREML, BOLT-REML, and RHE require individual data for target genotypes and phenotypes.

| Method | BOLT/GCTA-GREML, RHE | SUM-RHE | LDSC | SumHer |
| --- | --- | --- | --- | --- |
| Inputs | Individual genotype<br>Individual phenotype | Ref. trace<br>GWAS summary | Ref. LD scores<br>GWAS summary | Ref. SNP tag<br>GWAS summary |

**Figure 1:**
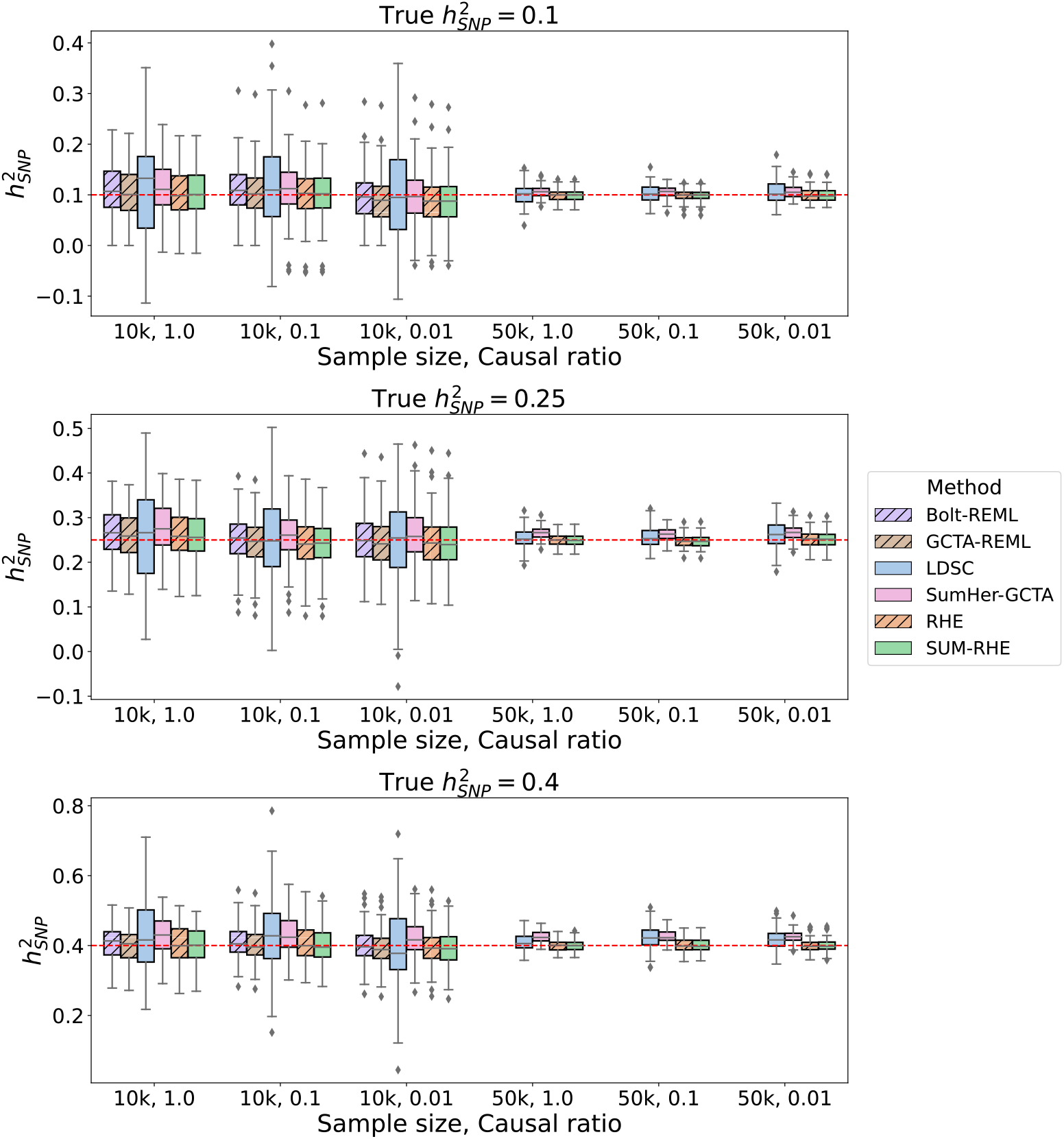
Comparison of SNP heritability estimates across methods. SUM-RHE heritability estimates are comparable to those from RHE or BOLT-REML/GCTA-GREML and are significantly more accurate than those of LDSC and SumHer. Due to computational limitations, BOLT-REML/GCTA-GREML was not run on the *N* = 50*k* simulations.

**Figure 2:**
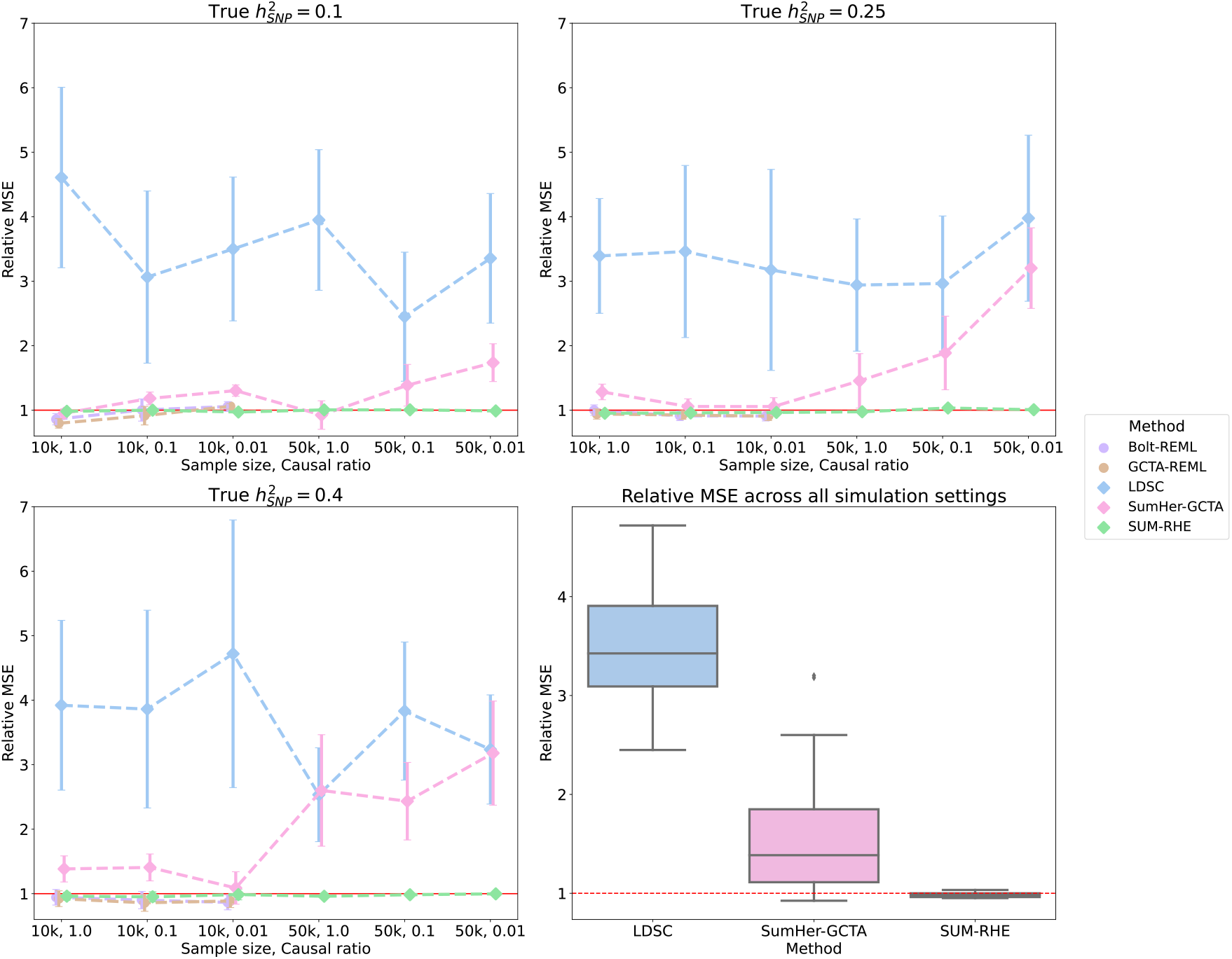
Mean Squared Error (MSE) of heritability estimates of each method relative to RHE. The dot and errorbar denote the relative MSE and the 95% CI calculated based on bootstrap resampling (using 10, 000 bootstrap samples), respectively. While the MSE of SUM-RHE is within *±*5% of the MSE of RHE, the MSE of LDSC ranges from 245% to 472% while the MSE of SumHer-GCTA ranges from 92% to 320%. BOLT-REML and GCTA-GREML have relative MSE in the range of 80% and 106% for the 10*k* samples.

SUM-RHE has substantially improved accuracy over other summary-statistic-based methods: LDSC exhibits MSE ranging from 244% to 478% relative to that of SUM-RHE (mean 356%), while SumHer-GCTA has MSE ranging from 94% to 331% relative to that of SUM-RHE (mean 167%). SumHer-GCTA has lower MSE than SUM-RHE for low heritability (*h*^2^ = 0.1) and high polygenicity (causal ratio *p* = 1.0). The improvement in MSE is particularly pronounced with smaller sample sizes (*N* = 10, 060).

We also tested the calibration of SUM-RHE by simulating phenotypes with 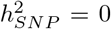 and testing the hypothesis of 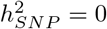 with rejection threshold *α* = 0.05 for both *N* = 10, 060 and *N* = 50, 112 to find that SUM-RHE is well-calibrated (Table 2).

**Table 2:**
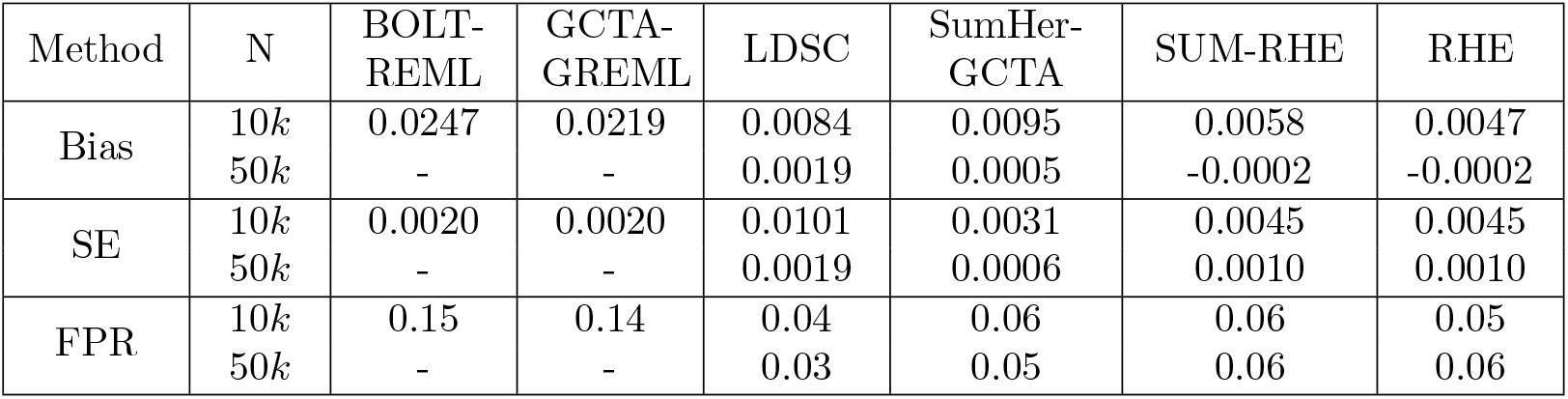
Calibration of the methods. We report the bias, SE, and the false positive rate (FPR) of each method in the setting where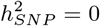. Due to computational limitations, GCTA-GREML and BOLT-REML were only run on 10*k* samples.

| Method | N | BOLT-REML | GCTA-GREML | LDSC | SumHer-GCTA | SUM-RHE | RHE |
| --- | --- | --- | --- | --- | --- | --- | --- |
| Bias | 10k | 0.0247 | 0.0219 | 0.0084 | 0.0095 | 0.0058 | 0.0047 |
|  | 50k | - | - | 0.0019 | 0.0005 | -0.0002 | -0.0002 |
| SE | 10k | 0.0020 | 0.0020 | 0.0101 | 0.0031 | 0.0045 | 0.0045 |
|  | 50k | - | - | 0.0019 | 0.0006 | 0.0010 | 0.0010 |
| FPR | 10k | 0.15 | 0.14 | 0.04 | 0.06 | 0.06 | 0.05 |
|  | 50k | - | - | 0.03 | 0.05 | 0.06 | 0.06 |

### 3.2 Simulations with a mixture model

We further test the robustness of our method by simulating phenotypes with a combination of large and small-effect SNPs. We selected the first *π* = 0.05 of the SNPs to account for *γ* = 0.25 of the total SNP heritability, while the rest of the SNPs accounted for the remainder. The causal SNPs were then selected at random with fixed probability *α*, such that the effect sizes for the large-effect SNPs were sampled from a different distribution than for the small-effect SNPs. Specifically, the effect sizes were sampled from two distributions:

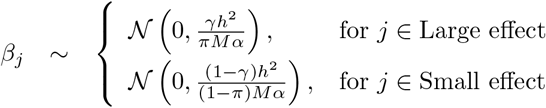

Figure 3 shows the boxplots of estimates from LDSC, SumHer-GCTA, SUM-RHE, and RHE, while Figure 4 plots the SE and MSE of the three summary-statistics methods (LDSC, SumHer-GCTA, SUM-RHE) relative to that of RHE. We observe that both LDSC and SumHer-GCTA show larger MSEs than SUM-RHE, similar to our previous simulations. LDSC has MSE in the range of 266% to 444% relative to that of SUM-RHE (mean 348%), and SumHer has MSE in 102% to 304% (mean 151%). These results indicate that SUM-RHE is robust under the mixture model simulations and attains accuracy comparable to individual-level methods across the scenarios we tested.

**Figure 3:**
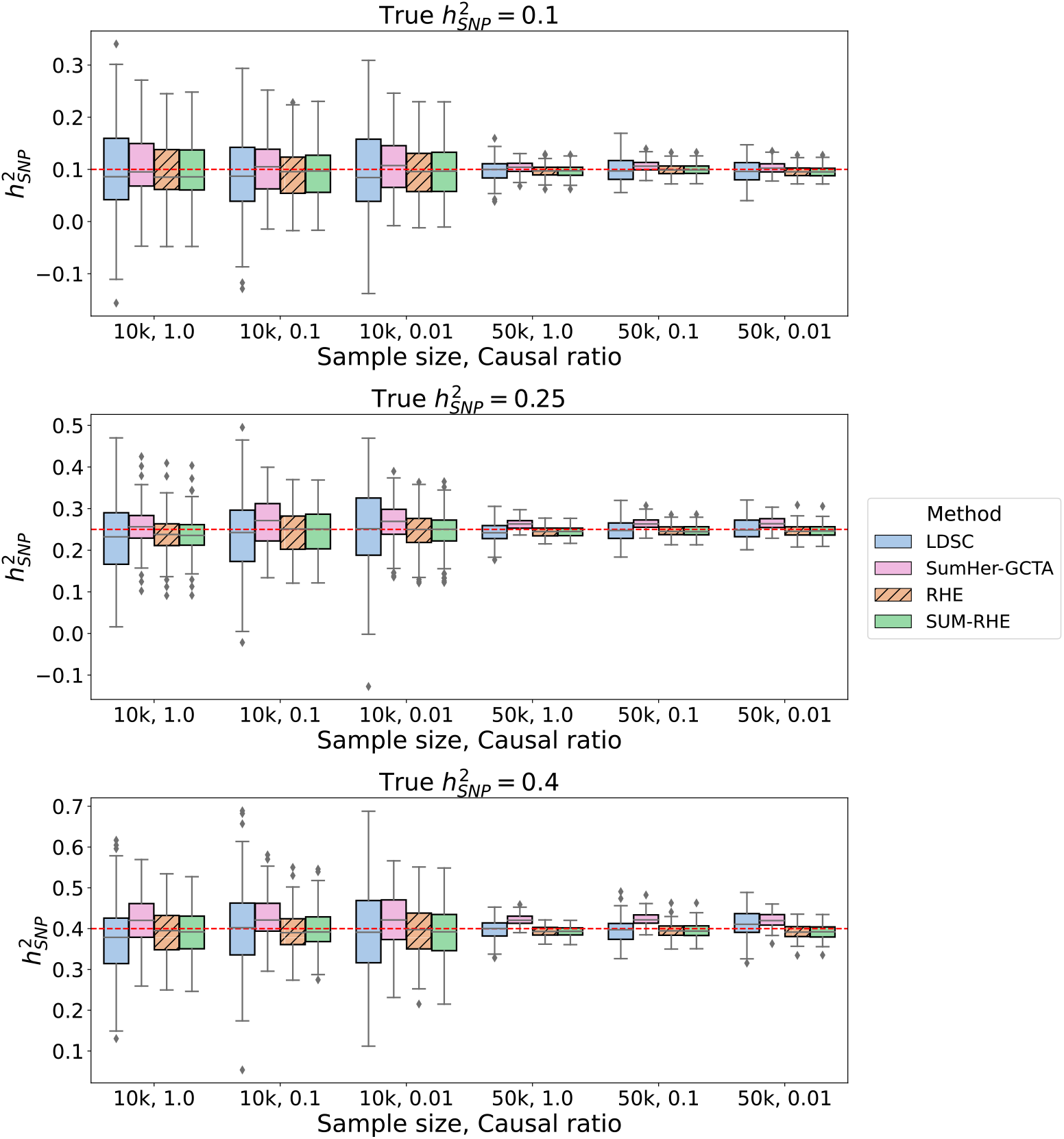
Comparison of SNP heritability estimates across methods on simulations with mixtures of large and small genetic effects.

**Figure 4:**
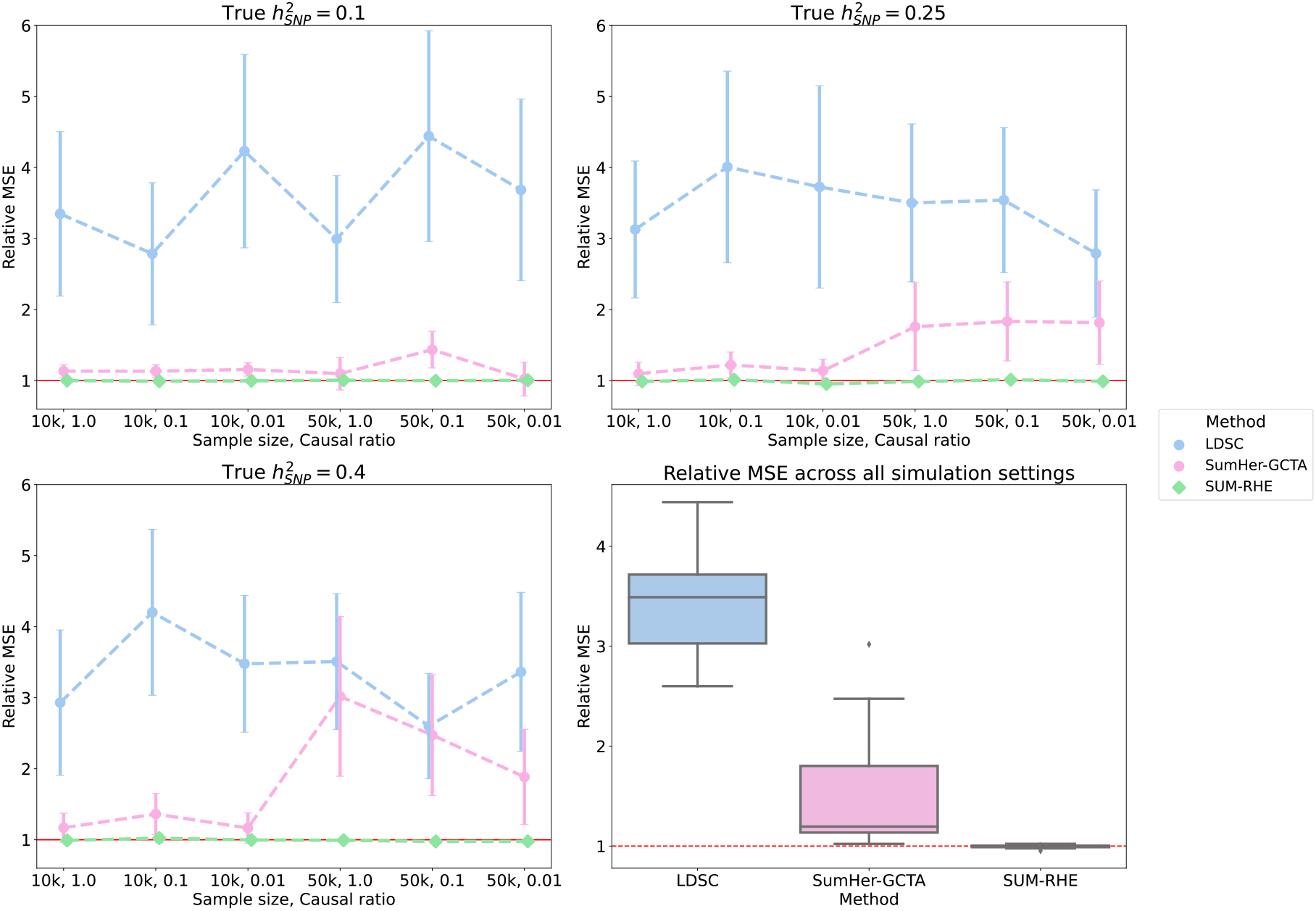
Comparison of MSE and SE of the summary statistics-based methods against RHE on simulations with mixtures of large and small genetic effects. Here we report the relative MSE of the three methods on the mixture-of-effects simulations. Their performances are similar to those in the previous simulations (Figure 2). SUM-RHE has an MSE within *±*5% relative to RHE.

**Figure 5:**
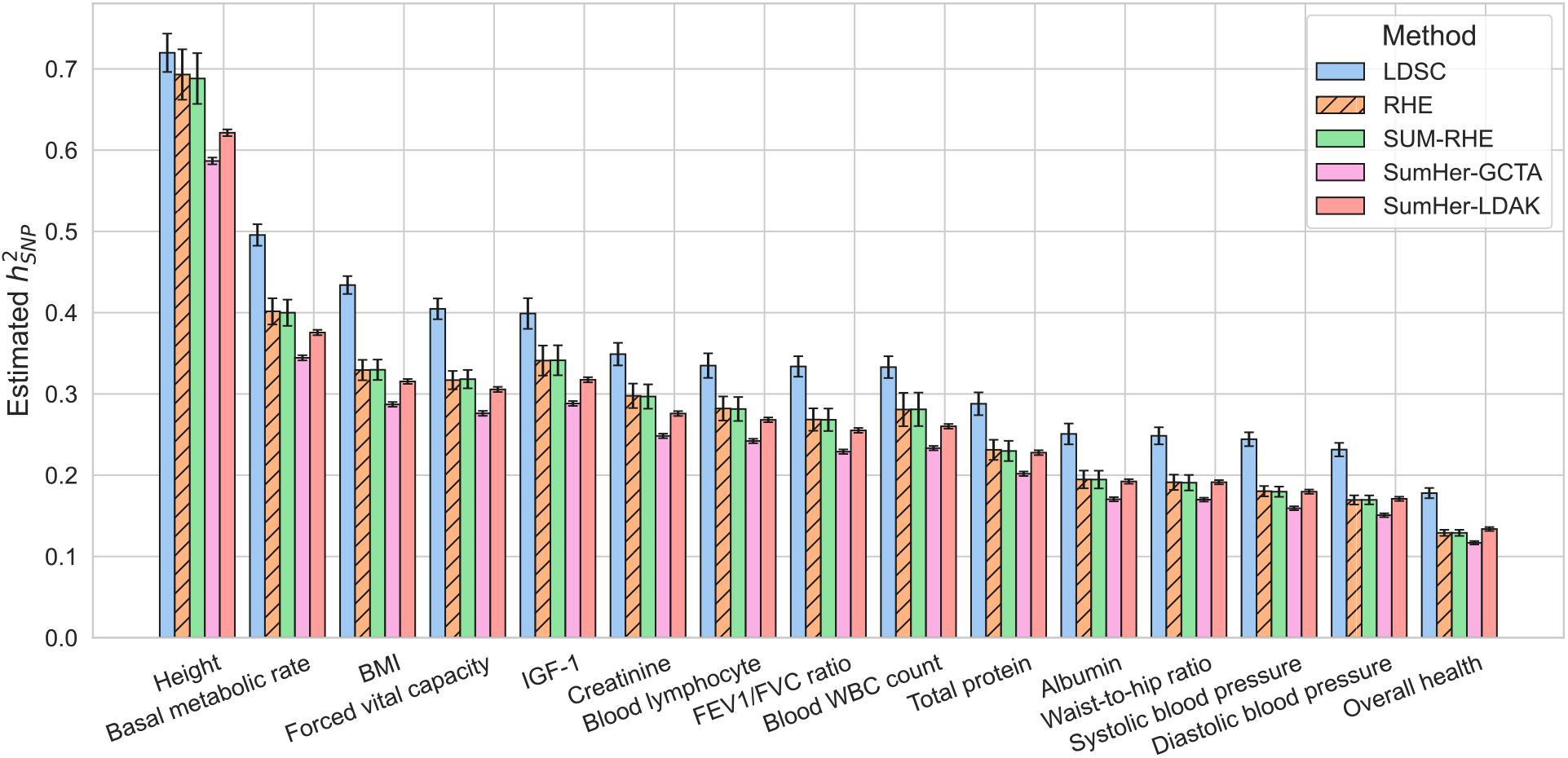
Application on real UK Biobank phenotypes.

### 3.3 Runtime measurements

We compared the runtime of SUM-RHE to other methods (Table 3). The heritability estimation step for all the summary statistic methods are computationally efficient irrespective of sample size. The generation of the reference statistics will depend on the size of the reference dataset but is typically a one-time computation that is relatively efficient even for datasets with hundreds of thousands of individuals with access to a compute cluster.

**Table 3:** Runtime estimates of the six methods. We ran each method on 10 replicates to measure wall clock time. For methods or tools that allow multithreading (BOLT/GCTA-GREML, SumHer, PLINK 2.0) we used 6 threads, run on the UCLA Hoffman2 computing nodes. SUM-RHE trace summaries were estimated by running the original RHE-mc codes (which does not support multithreading). PLINK 2.0 was used for calculating the GWAS summary statistics. All measurements are in seconds.

| Step | Sample size | BOLT-REML | GCTA-GREML | RHE | SUM-RHE | LDSC | SumHer |
| --- | --- | --- | --- | --- | --- | --- | --- |
| Reference statistic estimation | 281k | - | - | - | 14286.4 | 3120.8 | 697.3 |
| GWAS summary | 10k | - | - | - | 7.0 | 7.0 | 7.0 |
| Heritability estimation | 10k | 589.6 | 847.3 | 590.6 | 1.6 | 4.0 | 2.9 |
| Reference statistic estimation | 241k | - | - | - | 14686.0 | 2699.9 | 1180.7 |
| GWAS summary | 50k | - | - | - | 17.9 | 17.9 | 17.9 |
| Heritability estimation | 50k | - | - | 3032.6 | 1.3 | 4.0 | 1.6 |

### 3.4 Application to traits in the UK Biobank

Finally, we applied three of the summary statistic methods as well as one of the scalable individual genotype-based method (RHE) to real UK Biobank phenotypes measured on *N* = 291, 273 unrelated white British individuals paired with genotypes assayed on 454, 207 common array SNPs (we ran SumHer with both GCTA and LDAK SNP taggings for real phenotypes). Here we plot the 15 quantitative traits with the highest z-scores of SUM-RHE heritability estimates, from overall health (*z* = 34.3) to albumin (*z* = 17.9), ordered by the heritability estimates. For the summary statistics-based methods (LDSC, SUM-RHE, SumHer-LDAK/GCTA), we use in-sample statistics.

As expected SUM-RHE has estimates that agrees well with RHE estimates 5. SUM-RHE tend to lie in between those from LDSC and SumHer (with both GCTA and LDAK SNP taggings) consistent with our previous work ^11^.

## 4 Discussion

Here we propose a summary-statistics-based heritability estimation method, SUM-RHE, that has performance comparable to that of individual genotype-based methods. SUM-RHE is accurate, fast and highly portable. It uses a trace summary statistic calculated by aggregating stochastic trace estimates and PLINK GWAS statistics. In the era of large biobanks, SUM-RHE will be a useful tool in estimating heritability while maintaining the privacy of the patients.

We conclude with a discussion of limitations and directions for future work. First, heritability estimates from SUM-RHE are accurate under the assumption that the summary statistics are free of confounding due to population stratification and cryptic relatedness. In a setting where the summary statistics are affected by confounders, LDSC could potentially be more robust (as confounding would affect the intercept of LDSC while the slope would provide a robust estimator of heritability). Second, our preliminary experiments suggest that SUM-RHE retains its accuracy even when reference trace estimates are computed using a smaller number of random vectors on a smaller number of individuals (as low as *N*_*ref*_ = 30*k* and *B* = 1, 000 as seen in Figure S2). These results suggest that the computation of trace summaries can be even more efficient. Third, SUM-RHE is not applicable to the setting of MAF and LD-dependent architectures nor does it estimate partitioned or local heritability. These applications will require computation and release of partitioned trace summaries. We view this as a promising direction for future work.

## 5 Acknowledgments

This research was conducted using the UK Biobank Resource under application 33127. We thank the participants of UK Biobank for making this work possible.

