## Supplemental Figures for "Scalable summary statistics-based heritability estimation method with individual genotype level accuracy"

### Supplementary Information

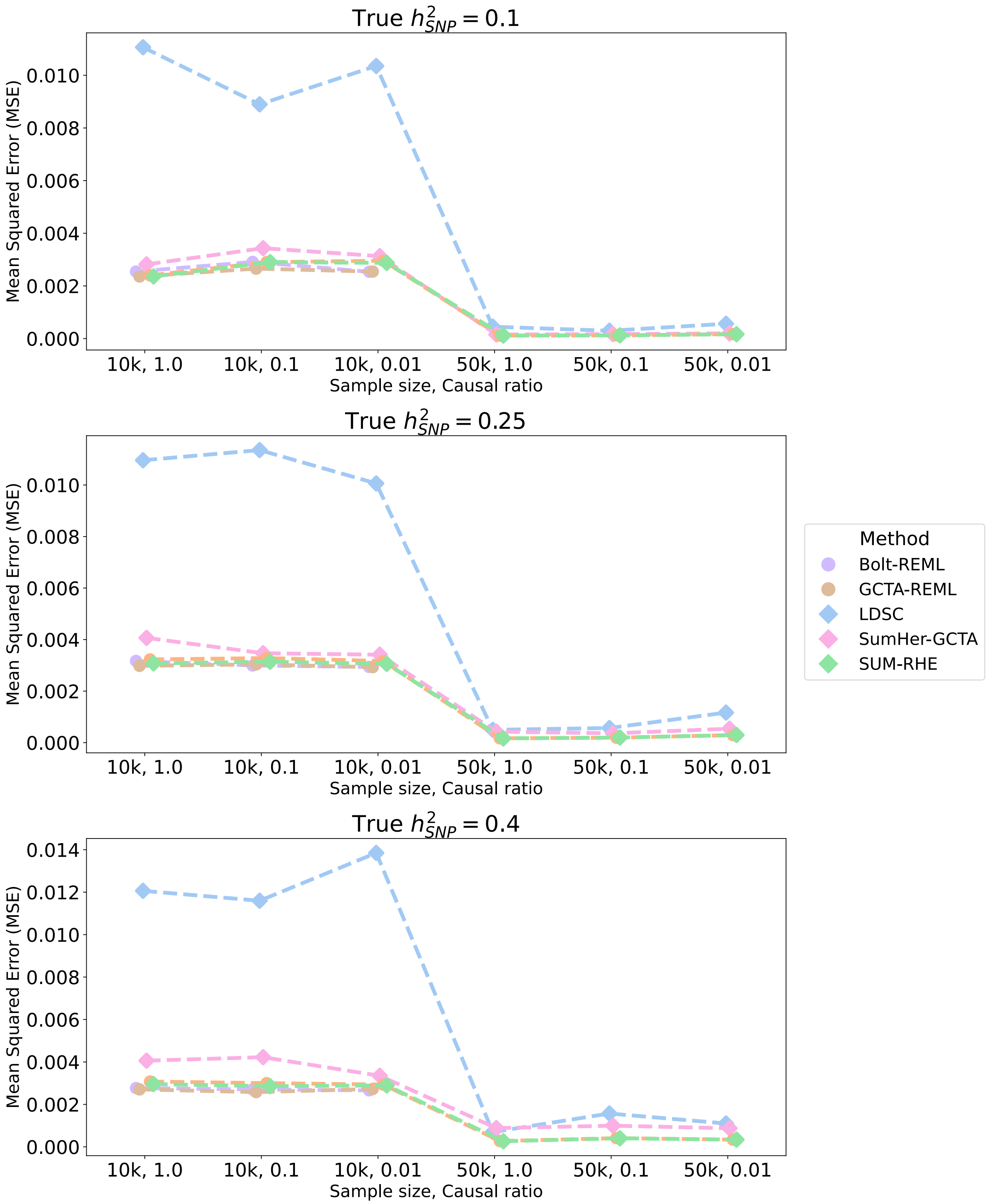


Figure S1: **Mean Squared Error (MSE) of heritability estimates of each method**

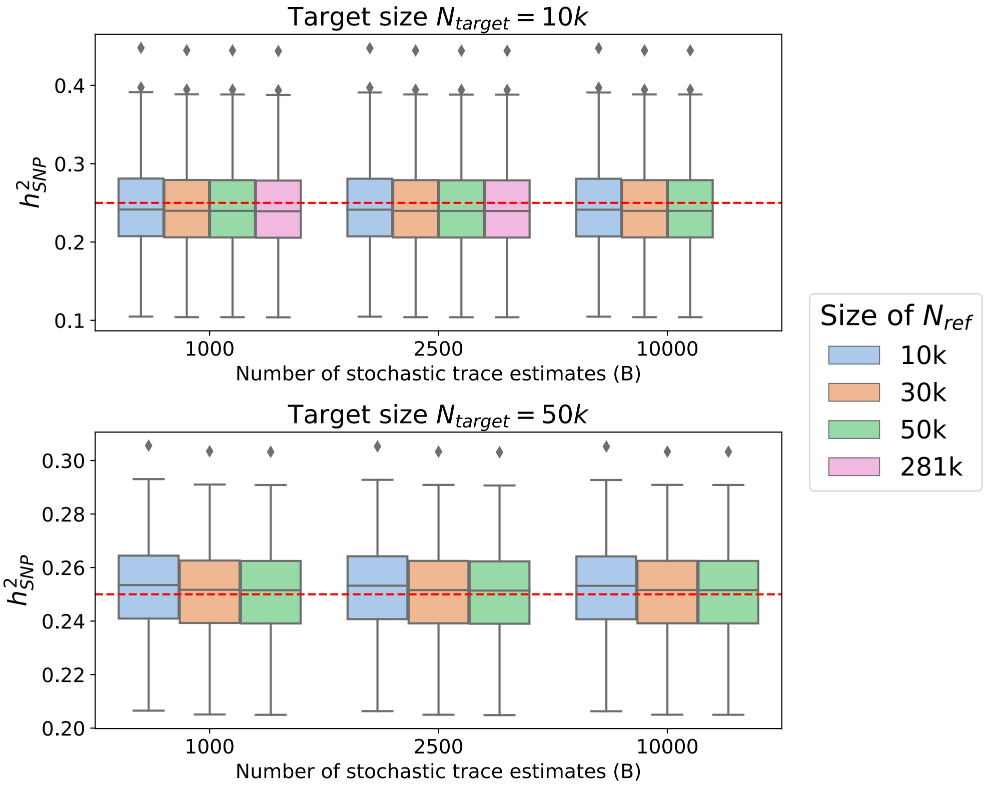


Figure S2: **Relationship between the trace estimates and accuracy of heritability estimates**. SUM-RHE requires the trace summary statistics as an input, which is converted from stochastic trace estimates on the reference dataset. Here we explored the impact of the size of the reference dataset ($N_{ref}$) and the number of stochastic trace estimates ($B$) from the reference dataset on the estimation of heritability in the target dataset. For the $N_{target} = 10k$ simulations, we generated a subset of the reference genotype with $N_{ref} = 281k$ to create $N_{ref} = 10k$, $30k$, $50k$ samples, from which we calculated trace summaries of varying number of runs. We then used these trace summaries to estimate heritability of $N_{target} = 10k$ simulation with true $h^2 = 0.25$ and causal ratio $p = 0.01$. We repeated this for the $N_{target} = 50k$ setting as well. Here we see that using a subset of size $N_{ref} = 50k$ and $B = 10,000$ gives trace estimates as accurate as using the entire $N_{ref} = 281k$ with $B = 2500$.
